# Label-free imaging of membrane potentials by intramembrane field modulation, assessed by Second Harmonic Generation Microscopy

**DOI:** 10.1101/2021.05.28.446105

**Authors:** Yovan de Coene, Stijn Jooken, Olivier Deschaume, Valerie Van Steenbergen, Pieter Vanden Berghe, Chris Van den Haute, Veerle Baekelandt, Geert Callewaert, Stijn Van Cleuvenbergen, Thierry Verbiest, Carmen Bartic, Koen Clays

## Abstract

Optical interrogation of cellular electrical activity has proven itself essential for understanding cellular function and communication in complex networks. Voltage-sensitive dyes are important tools for assessing excitability but these highly lipophilic sensors may affect cellular function. Label-free techniques offer a major advantage as they eliminate the need for these external probes. In this work, we show that endogenous second harmonic generation (SHG) from live cells is highly sensitive to changes in membrane potential. Simultaneous electrophysiological control of a living (HEK293T) cell, through whole-cell voltage clamp reveals a linear relation between the SHG intensity and membrane voltage. Our results suggest that due to the high ionic strengths and fast optical response of biofluids, membrane hydration is not the main contributor to the observed field sensitivity. We further provide a conceptual framework that indicates that the SHG voltage sensitivity reflects the electric field within the biological asymmetric lipid bilayer owing to a nonzero 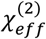 tensor. Changing the membrane potential without surface modifications such as electrolyte screening offers high optical sensitivity to membrane voltage (~40% per 100 mV), indicating the power of SHG for label-free read-out. These results hold promise for the design of a non-invasive label-free read-out tool for electrogenic cells.

## Introduction

Strict control over the ionic imbalance across the plasma membrane is of crucial importance for cellular function and communication. During a neuronal action potential, the ionic imbalance between the intra- and extracellular fluids results in an electrochemical potential difference of up to 100 mV. Considering that this voltage drop occurs over a bilayer of phospholipid molecules, typically only a few nanometres wide, enormous electric field strengths in the order of 100 kV/cm, are attained.^1^ During neuronal firing, these membrane-spanning DC-fields are modulated at kHz frequencies and therefore require a read-out tool with a millisecond temporal resolution.^2^ These immense, spatially confined, electric fields in biological structures can effectively couple with light through various physical mechanisms, and can therefore be used for a non-invasive optical interrogation of cellular activity. Therefore, electro-optical techniques are regarded as a very promising platform, continuously leading to novel insights in neuroscience. ^2^

Potentiometric membrane dyes, which report through a modulation of their fluorescence intensity, are valuable by their relative ease of use as compared to microelectrode recordings. Two-photon fluorescence (TPF), a nonlinear optical (NLO) effect where two near-infrared (NIR) photons simultaneously combine to excite a fluorophore in the UV-blue range, can greatly enhance light penetration into tissue as the excitation wavelength is situated within the spectral window of biological tissue transparency (700 to 900 nm).^3^ Yet, the associated excited states limit experimental duration due to probe photobleaching. An interesting alternative which does not rely on excited states, is provided by another NLO effect called second harmonic generation (SHG). In SHG, two photons combine through virtual states to scatter one photon at twice the excitation frequency. SHG is a second-order nonlinear optical (SONLO) process and, as all even-order NLO processes, only susceptible to non-centrosymmetric materials, both at the molecular and supra-molecular level. The electric field dependence of SHG is governed by a third-order nonlinear optical (TONLO) response, with a sensitivity greatly exceeding that of fluorescence.^4,5^ SHG voltage imaging has been primarily based on engineered molecules with large hyperpolarizabilities and amphiphilic properties,^6–9^ with new, unlabelled approaches now beginning to emerge. ^10–12^

All techniques using membrane dyes can interfere with the delicate cell membrane as they often have significant dipole moments leading to local field disruption and alteration in membrane capacitance, thereby affecting both action potential conductance and membrane transport.^2,13^ Moreover, optical labels were found to significantly modify flip flop dynamics relative to unlabelled systems.^14^ Therefore, label-free read-out of membrane potential, using SHG offers an attractive alternative for imaging living cells.

Mammalian cells are ideal candidates for label-free SHG imaging given the structure and electrical properties of their membranes. The plasma membrane and associated diffuse layers act as capacitors where spatially confined potential differences can be attained, while the interfacial region between solid (lipid bilayer) and liquid (electrolyte solution) imposes a nonlinearity through a local change in refractive index. The electric potential across the cell membrane follows a complex profile, changing in magnitude and sign over sub nanometre length scales. These changes occur within the coherence length of optical microscopy and therefore require detailed knowledge of the interfacial effects residing at the cell membrane. An intricate electric potential profile arises from the combined effect of the lipidic orientation, the active maintenance of an ionic imbalance and an overall negative surface charge (see Figure 1).

**Figure 1:**
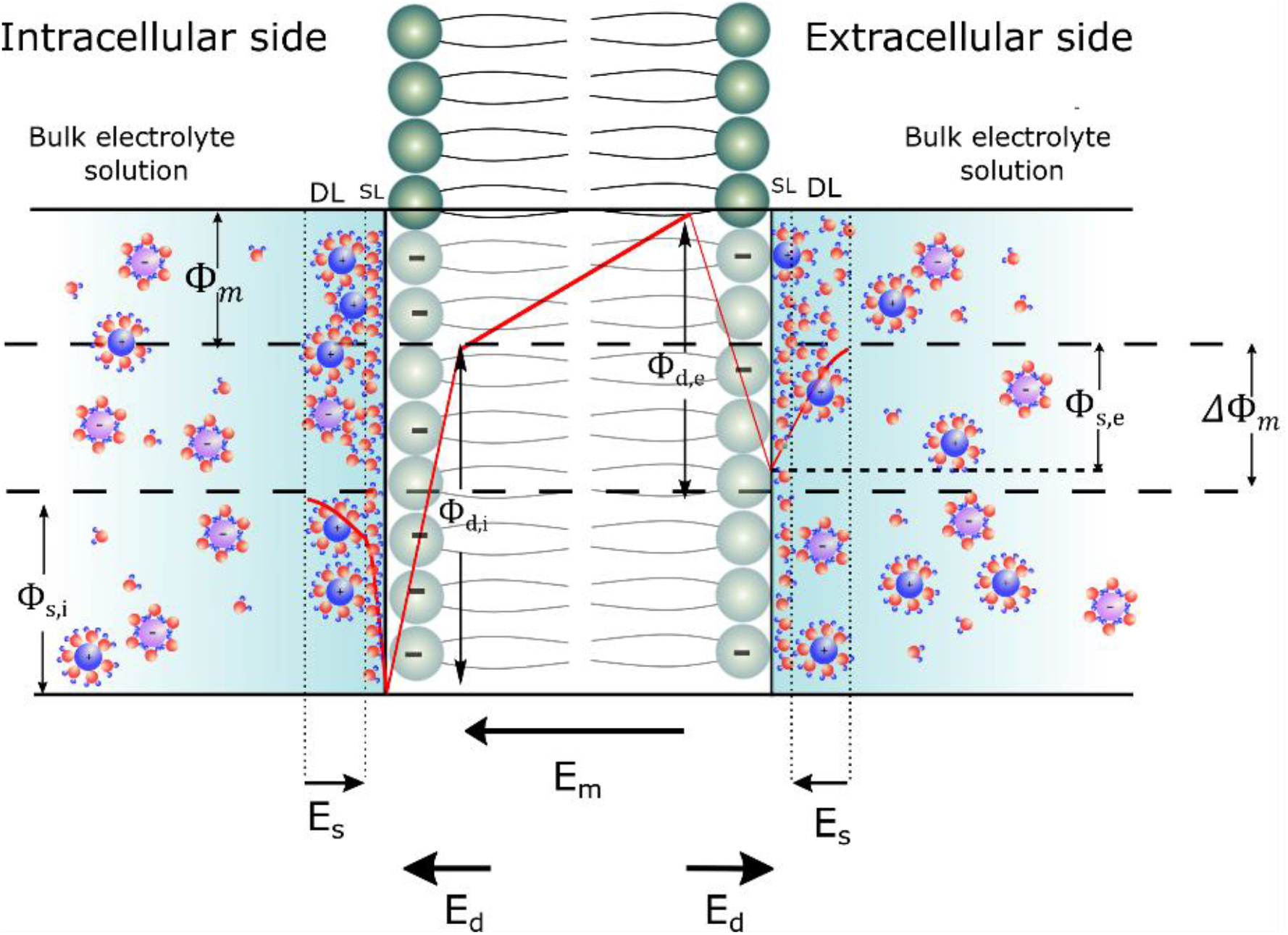
Schematic representation of the different electric potentials (red lines) which contribute to the total membrane potential (TMP), **ΔΦ_m_** (between the dashed lines) and the corresponding electric fields (black arrows) along the plasma membrane. DL: diffuse layer, SL: Stern layer. The total TMP, **ΔΦ_m_**, arises from three major components: (1) the intramembrane potential (IMP), **Φ_m_**, (2) the dipole potential, **Φ_d_**, and (3) the surface potential, **Φ_s_**. **Φ_m_** merely results from an ionic imbalance across the lipid bilayer, that is actively maintained by live cells. Due to an asymmetrical lipid distribution between the inner and outer leaflet,^15^-^17^ the inner leaflet is more negatively charged and hence has a larger surface potential compared to the luminal side. The sign of the resulting electric field vector is equal to that of **Φ_m_** in a cell at rest (pointing inwards, towards the cell centre). The dipole potential **Φ_d_** arises from the unfavourable parallel alignment of phospholipids and water dipoles, and is manifested between the hydrocarbon interior of the membrane up to the first few water layers adjacent to the lipid head groups. **Φ_d_** is unaffected by changes in the ionic composition of the solutions ^18^ and depends entirely on the packing density, cholesterol content, ^19^ and membrane curvature. ^20,21^

For a charged surface in contact with water, SHG mainly originates from the interfacial water molecules.^22^ Both the inner and outer membrane interfaces have second- and third-order susceptibility tensors (χ^(2)^ and *χ*^(3)^) that originate from the Stern and diffuse layers, respectively.^23–25^ *χ*^(2)^ reports on symmetry breaking at the interface and neatly aligned water molecules, ^26,27^ while *χ*^(3)^ finds its origin in the layer of water molecules, regardless of their orientation, extending into the aqueous phase according to the Debye length, which is dependent on the ionic strength.^28^ Very recently, it was shown that K^+^-induced depolarization of neurons could be monitored label-free,^12^ with an SHG source being attributed from the orientation of interfacial water molecules,^22^ and a frequency doubling efficiency directly relating to the transmembrane potential (TMP).^29^ However, this system reflects a particular situation, where changing the ionic strength on only one side modifies the electric potential profile locally due to charge build up around ion channels on a slow temporal scale (seconds).^11^ This is in contrast to a physiologically relevant system that exhibits millisecond transport of ions.

It is well documented, by the pioneering work of the group of K.B. Eisenthal, that a *χ*^(3)^ dependent SHG signal caused by interfacial water is in direct correlation with the thickness of the Debye length at the interface.^22^ Large Debye lengths hence induce more efficient SHG responses of interfacial water.^25^ The quadratic response of the SHG intensity as a function of membrane voltage can be explained in that case by a long decay of the surface potential into the bulk and hence a large *χ*^(3)^ dependent response and is a hallmark of electric field induced SHG (EFISH). This is well in line with the fact that the only report of label-free voltage (based on electrode control) read-out of a bilayer using SHG was observed in an artificial bilayer system having an ionic strength of around 100 μM which is about 1000 times lower than in biological fluids. ^10^ For low ionic strengths the Debye length can be several hundreds of nanometres while the high ionic strengths in biofluids (>100 mM) induces a sub-nanometre Debye length and causes the *χ*^(3)^ term from interfacial water to lose its dominance. ^25,30^ Therefore, dynamically tracking the slow changes^11,12^ in surface potential using SHG does not seem convenient for monitoring physiological membrane voltage changes. Rather than mapping the change in surface potential through a change in *χ*^(3)^, our data shows that it is possible to map the change in effective *χ*^(2)^ of the lipid bilayer itself, without interfering with the surface potential. Moreover, the change in SHG efficiency, based on the change in intramembrane potential of a biological cell membrane is found to be superior in terms of response and voltage sensitivity.

## Results

Human embryonic kidney 293T (HEK293T) were chosen for our experiments. They are a popular first choice in a wide range of scientific disciplines due to their easy maintenance, robustness and their ease of transfection. HEK293T cells were subjected to SHG imaging at a laser excitation wavelength of 900 nm while the TMP was held at different voltages using the whole-cell patch-clamp technique. SHG was detected in the forward direction by a narrow band-pass filter at 450 nm while TPF was detected in the backward direction using a 470 nm long-pass filter (Figure 2a). Acquisition was performed using the photon counting mode of the photomultiplier tubes in order to increase the signal to noise ratio. A field of view of 200 x 200 μm at a resolution of 640 x 640 pixels was scanned to localize the tip of the patch pipette and the cell to be studied (Figure 2b). Once the cell was identified, a line along the top or bottom of the plasma membrane perimeter was selected where the incoming light polarization has the same orientation as the electric field over the membrane and the highest signal is expected (Figure 2b and c). The region of the patch pipette was always excluded. A pixel dwell time between 100 and 500 μs was used, depending on the laser power (20-40 mW at the sample), and recordings were performed for up to 100 seconds. Each line scan over the entire region of interest is plotted as a function of time in Figure 2c. Figure 2d shows the voltage applied across the cell membrane and the recorded SHG and TPF intensities as a function of time. Whereas the TPF signal did not change appreciably when the membrane voltage was stepped between −100, 0 and +100 mV, the SHG intensity clearly decreased following membrane depolarization.

**Figure 2:**
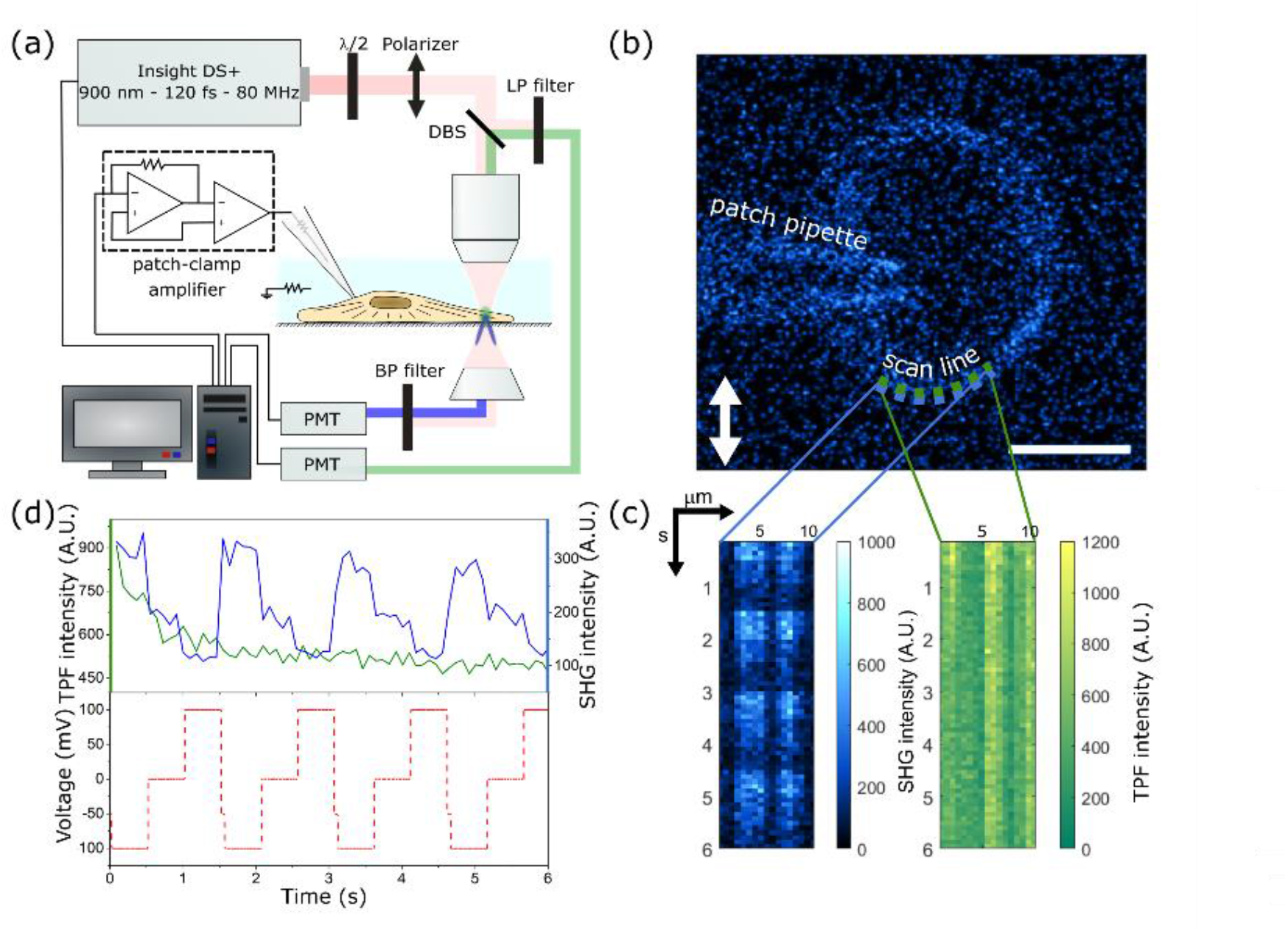
Scheme of the experimental design. (a) Multiphoton/ patch-clamp set-up allowing simultaneous imaging of TPF (reflection, green) and SHG (transmission, blue) while whole-cell voltage clamping. (b) SHG image of the patched cell. Scale bar is 10 μm, and the white arrow indicates the incoming light polarization direction. The dashed lines depict the region of interest at the membrane where the SHG (blue) and TPF (green) intensities were measured simultaneously, as a function of membrane voltage. A Matlab script was written in order to represent the data as in panel (c) using Gaussian smoothing, showing the line scans of SHG (left) and TPF (right) intensities where the y-axis indicates the time and the x-axis the distance along the cell membrane. Panel (d) shows the average SHG (blue trace) and TPF (green trace) intensities along each line as a function of time, and the imposed voltage protocol (red trace).

The SHG intensity changed with the applied electric field only when the seal resistance before rupturing the cell membrane was sufficiently high (> 800 MΩ). It was also observed that as leak current increased (Figure 3c) over time, the peak intensity of the SHG signal concomitantly decreased (Figure 3a) while the TPF signal recorded in reflection mode only displayed an initial bleaching phase (Figure 3b). We do not have access to a spectrometer-coupled detection scheme. However, given the fact that the TPF shows no correlation with the applied voltage, any fluorescence passing the SHG filter would result in an offset of the absolute intensity, while the sensitivity would remain unaffected. The increase in leak current over time is likely related to a decrease in seal stability and/or a partial membrane electroporation caused by the applied 100 mV voltage pulses.

**Figure 3:**
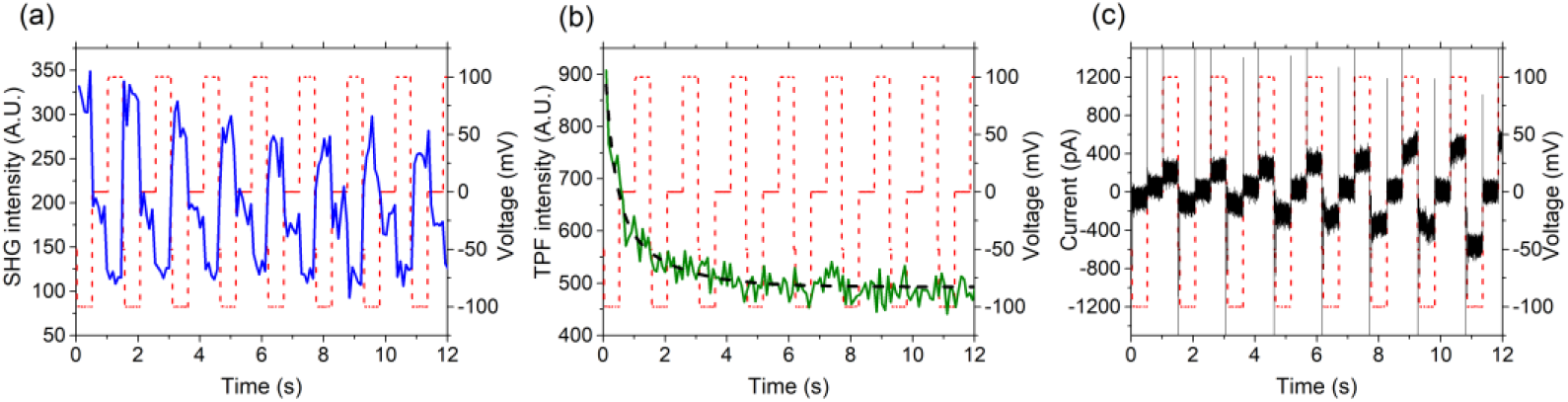
SHG intensity (a – blue trace), TPF intensity (b – green trace) and membrane current (c – black trace) recorded as a function of time during consecutive voltage steps (red traces superimposed). The superimposed dashed black trace in (b) is a bi-exponential fit to indicate photobleaching. The membrane potential was intermittently stepped from the holding potential (−50 mV) for 500 ms to −100 mV followed by a 500 ms step to 0 mV and a 500 ms step to +100 mV before stepping back to holding potential for 16 ms. Pixel dwell time was 200 μs and pixel size was 0.5 μm.

The latter is supported by the finding that a voltage-clamp protocol using smaller voltage steps between −50 and +50 mV generated a more steady SHG signal (Figure 4). In addition and in line with previous reports, ^10–12^ spatial heterogeneity in SHG intensity and sensitivity likely reflecting local changes in membrane curvature and heterogeneities in lipid composition could be observed.

**Figure 4:**
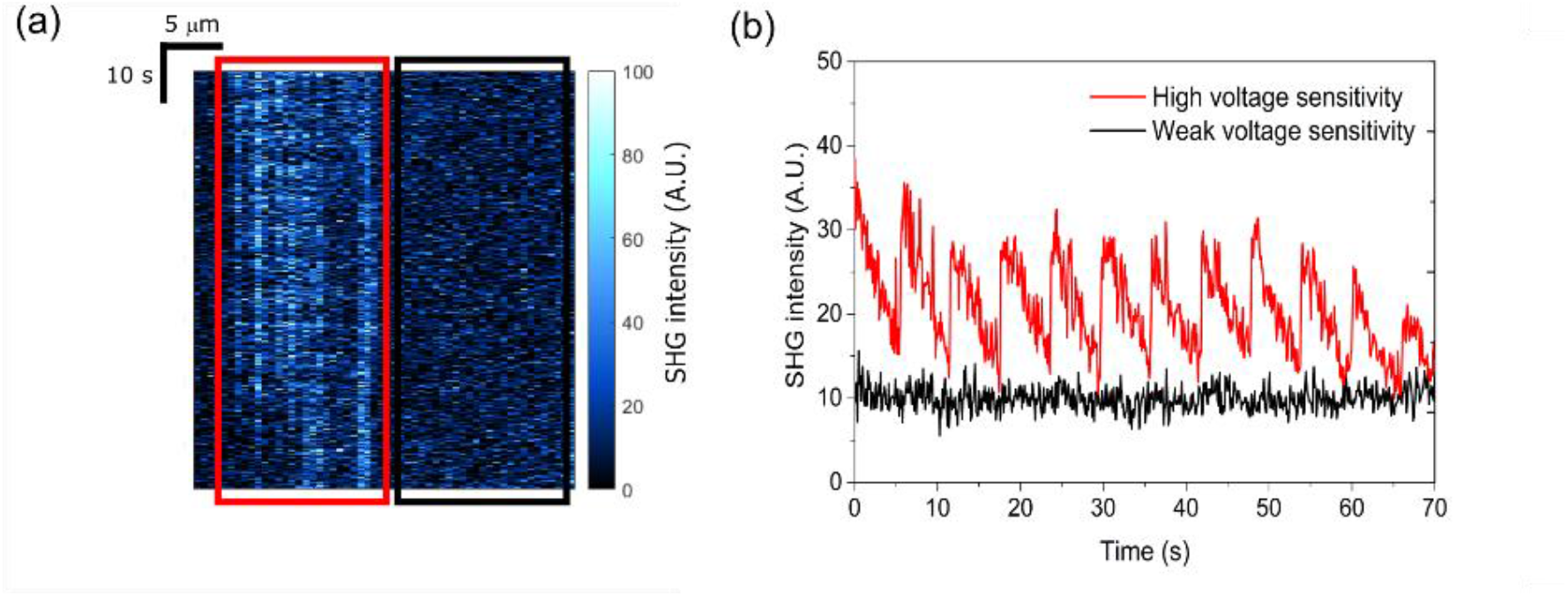
(a) Raw SHG intensity for each line scan along the cell membrane for a stepwise patch clamp protocol from −50 to 50 mV in 10 mV increments at 500 ms per increment. After each increment, the holding potential was held at −50 mV for 16 ms. The red box shows the area along a cell membrane region with high voltage sensitivity (corresponding to the red curve in (b)), while the black box shows an inactive region (corresponding to the black curve in (b)).

Data collected from 3 different cells using identical microscope alignment, settings and laser power were used to calculate the mean SHG voltage response. The voltage sensitivity was calibrated by measuring the SHG intensity while changing the applied membrane voltage from −50 to +50 mV in 10 mV increments with 500 ms per step. The relative intensities were calculated by taking the difference between the SHG intensity at the holding potential of −50 mV and at 0 mV, divided by the SHG intensity at 0 mV. The relative mean SHG intensity was 38.9 +- 1.7 % per 100 mV (Figure 5). It has to be noted no SHG was detected near the pipette opening. Moreover, no SHG sensitivity to the applied voltage was observed from the capillary walls, as the applied potential is too low in order to detect this recently reported effect.^31^

**Figure 5:**
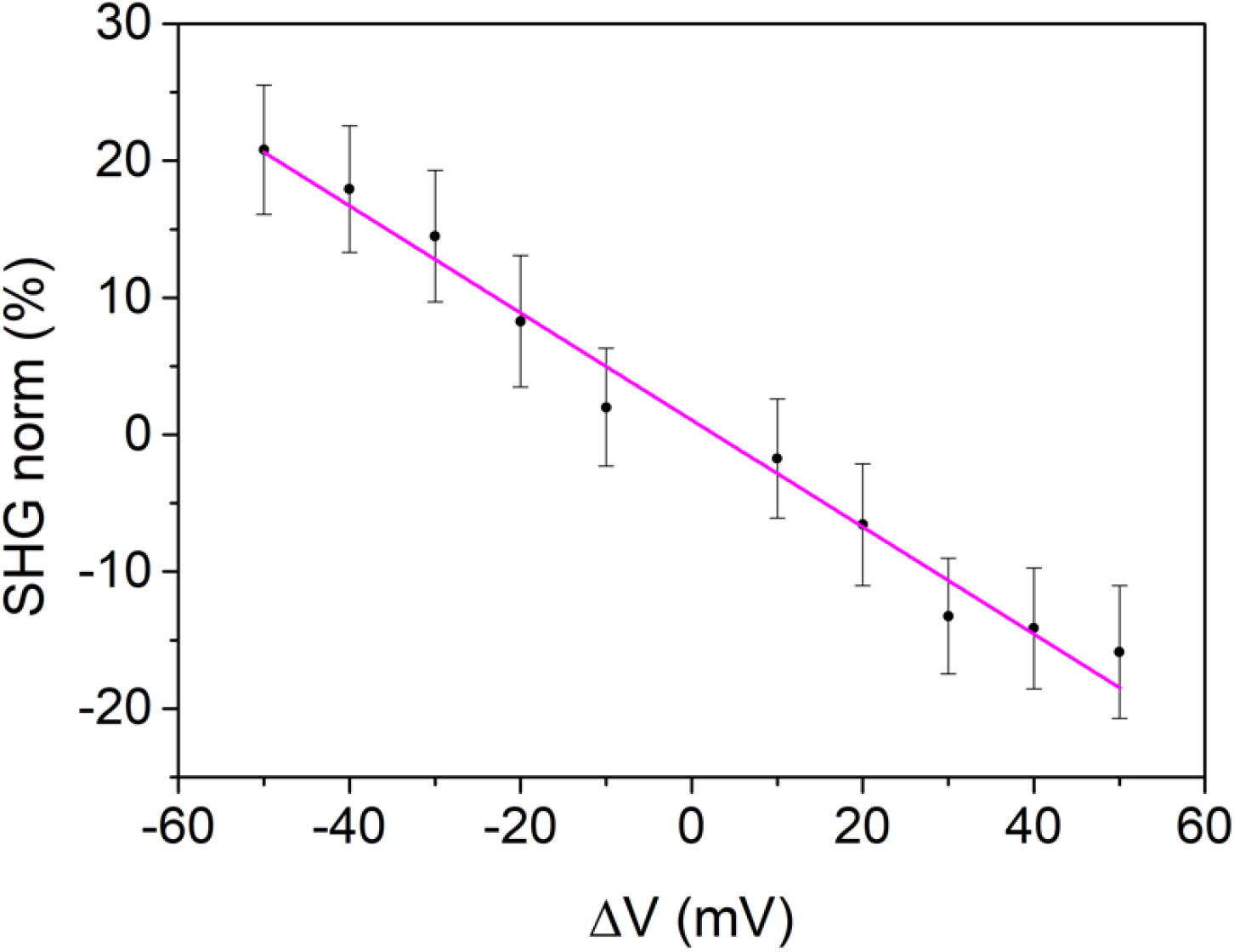
Plot of relative SHG intensity changes (expressed in percent) as a function of membrane voltage. The magenta line shows a linear regression of the averaged SHG intensities. Data based on 19 voltage sweeps in 3 different cells.

## Discussion

The TMP (Δ**Φ_m_**), gauged or controlled by voltage clamp, is blind in differentiating between SP (**Φ_s_**) and the IMP (**Φ_m_**), and also contains no information about the local electric field strength. In whole-cell patch clamping the electric field component resulting from the IMP (**Φ_m_**) is predominantly influenced. For high ionic strengths, such as in biofluids, the cell membrane can be regarded as three capacitors in series.^20^ Two capacitors are associated with the diffuse layers (inside and outside), while the third one represents the lipid bilayer. An electric potential drops most over the smallest in series capacitor. Given that the permittivity of water in the diffuse layer (ε ~ 80)^32^ is much larger than that of the lipid bilayer (ε ~ 2), the screening length at the surface of a cell approaching zero, and therefore the electrical potential difference resulting from an ionic imbalance or an externally applied electric field drops exclusively within the membrane fatty interior. ^33^ For a biological cell membrane, this capacitance ratio (taking into account the permittivity, the Debye length and the membrane thickness) is in the order of 1% and entails that the gross of the potential drop is generated inside the hydrophobic core of the membrane.^34^ Therefore we can relate *Δ***Φ_m_**, applied by voltage clamping to **Φ_m_** inside the lipid bilayer. Moreover, since the thickness of the bilayer does not change during measurements one can directly relate changes in **Φ_m_** to changes in *E_m_*. Since the electric field profile associated with the cell membrane extends only on a length scale of nanometres, we can neglect quadrupolar effects at NIR excitation wavelengths. Therefore, the electric field, oscillating at twice the excitation frequency (*E_2ω_*) can be related to the induced second-order nonlinear polarization at 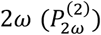 and written as:

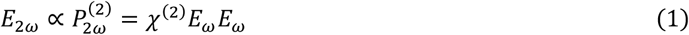

where *χ*^(2)^ is the second-order susceptibility tensor, and *E_ω_* the electric field of the fundamental light. In the presence of a third electric field at zero frequency, a third-order nonlinear induced polarization, oscillating at 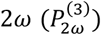 has to be considered:

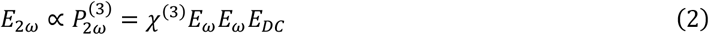

where *χ*^(3)^ is the third-order nonlinear susceptibility tensor and *E_DC_* the zero-frequency electric field affecting the material. The following equations will specifically apply to a system where: (1) the ionic strength is high (> 0.1 M), (2) in the absence of phase interference (i.e. the entire electrostatic profile decays over only 10 nm), (3) for SHG transmission geometries and (4) off-resonance. Hence, an effective susceptibility tensor, 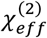, can be expressed as the combination of SONLO and TONLO effects leading to a DC-field dependent SHG efficiency:

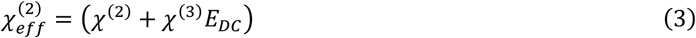

The induced electric field oscillating at 2ω can be written as the combination of equations (1) and (2) and can be related to the SHG intensity as:

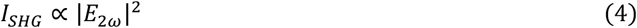

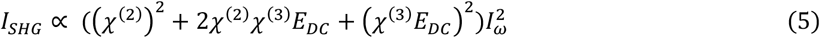

The contribution of the phospholipids themselves, which are non-centrosymmetric, is usually neglected since the second-order susceptibility of the bilayer is resulting from the difference in *χ*^(2)^ related to the difference in surface charge.^29^ However, given the asymmetric nature in lipid distribution, we propose that phospholipids should be accounted for, through an effective susceptibility of the bilayer itself 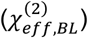. Therefore, the SHG at the membrane region for a living cell can be written as a modification of equation 4:

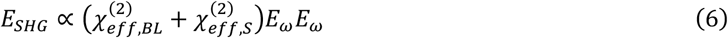

The effective susceptibilities describe the contribution of SONLO and TONLO induced polarization within the lipid bilayer 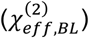 and at the surface 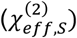 to the total SHG. When the membrane potential is effectively controlled by voltage clamp, we can separate a static and a dynamic contribution, due to the fact that the voltage modulation occurs entirely over the lipid bilayer, to the field at 2ω:

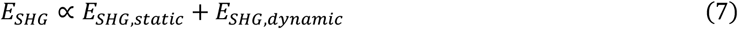

where *E_SHG,dynamic_* is the SHG component depending on the TMP controlled by voltage clamp. As mentioned before, the SP is not significantly affected by changing the TMP by voltage clamp, due to the relative high permittivity of interfacial water. Therefore, the static component can be expressed as:

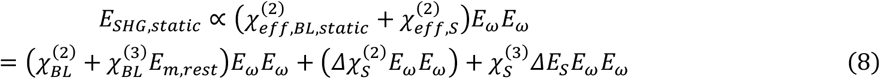

with *E_m,rest_* the membrane electric field when the cell is at its resting membrane potential, with 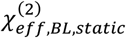 the effective susceptibility at that particular membrane potential. The magnitude of the latter is determined by the molecular NLO polarizabilities of the lipids, lipid content, the asymmetry in lipid content and type, and the magnitude of *E_m,rest_*. For a charged interface in contact with a liquid phase, H_2_O has two molecular second hyperpolarizabilities expressed as the sum of both the electronic (*γ_e_*) and rotational (*γ_r_*) contributions, the latter being dependent on the orienting static electric field. The interfacial water (IF) contribution at the molecular level, can be written as the effective surface susceptibility contribution on each side of the membrane:

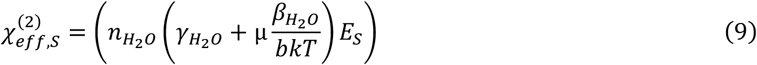

with *n*_*H*_2_*O*_ the number density of water molecules. *β*_*H*_2_*O*_ and *γ*_*H*_2_*O*_ are the molecular first and second hyperpolarizabilities of water respectively, whereas μ is the permanent dipole moment of water. kT accounts for the thermal energy and *E_s_* represents the surface electric field. A constant *b* is introduced that accounts for the susceptibility elements within the experimental system. It is obvious that the number of water molecules contributing to this response, depending on the Debye length, is an important factor in the equation. In a live cell, short Debye lengths due to high ionic concentrations would imply a relatively small influence of interfacial water. Equation 9 only describes one interface, in reality there are two, with electric fields opposite in sign (see Figure 1). For the lipid bilayer, only the electronic part of *γ* will matter as no significant electric field induced orientation occurs.^33^ Consequently, we can write the dynamic component, under the control of voltage clamp as:

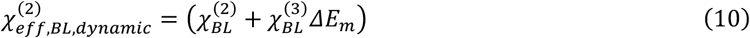

with *ΔE_m_* the difference between the membrane electric fields resulting from the resting membrane potential of the cell and the potential applied by the voltage clamp potential. Assuming the same laboratory frame for the entire interfacial region, the oppositely oriented phospholipids result in a change in sign of *β_zzz_* in each layer, (as opposed to *γ_zzzz_*). For a given TMP, the electric field along the phospholipid bilayer (*E_m_*) resulting from **Φ_m_**, is constant and does not change sign (within the bilayer itself), meaning that total destructive interference is avoided only if the number density and/or the *χ*^(2)^ differ between both leaflets. Lipid packing density and hydration also differs between the inside and the outside due to geometrical constraints and different headgroup interactions.^35,36^ Since the lipid bilayer consists of an asymmetrical distribution (in terms of phospholipid species and number density) of non-centrosymmetric opposing molecules, 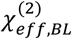 will be nonzero. We have to stress that, although SHG occurs as a complex interaction between three main electric fields (two surface electric fields and one intramembrane electric field), the SHG has multiple sources (membrane lipids, interfacial water, and possibly membrane proteins), each within their molecular frame, occurring in the same laboratory frame. Considering a unidirectional, constant DC-field (*E_m_*) within the lipid bilayer, the SHG field can be written as:

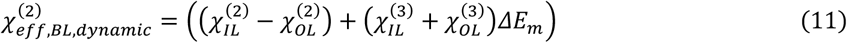

and the total SHG intensity coming from the membrane region from a voltage clamped cell as:

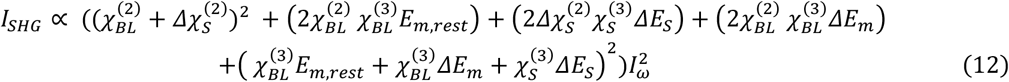

with 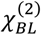 and 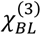 the second- and third-order susceptibility tensors of the lipid bilayer, 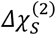 represents the net second-order susceptibility tensor of the interfacial water due to the boundary potentials with opposite signs at both sides of the bilayer. 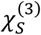 is the third-order susceptibility tensor arising from bulk water molecules where the SP decays over. The net effect of the third-order response depends on the residual potential of the SPs, which are opposite in sign at both sides of the membrane, merely due to the opposite sign in surface electric field (see Figure 1).

The voltage sensitive SHG signal in HEK293T cells can thus be expressed as:

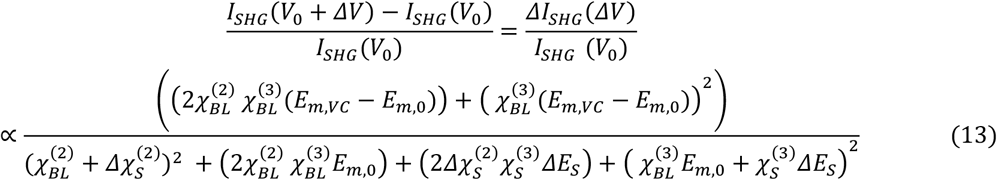

with *I_SHG_*(*V*_0_) and *I_SHC_*(*ΔV*) the SHG intensity at 0 mV and the clamped potential respectively. Although we normalize the SHG intensity by its intensity measured at 0 mV (*Δ***Φ_m_** = 0), it will result in a nonzero IMP (**Φ_m_** ≠ 0), because of the previously discussed membrane asymmetry. We termed the intramembrane electric field resulting from this unknown but nonzero IMP at 0 mV ‘*E*_*m*,0_’, while the electric field related to the potential applied by voltage clamp is termed ‘*E_m,VC_* The dynamic component in equation 13 is thus limited to the terms which contain the second- and third-order susceptibility tensors of the lipid bilayer itself, as the induced voltage bias will exclusively drop over the central part of the bilayer.

Our results indicate a dominating linear response which, by looking at equation 13, is likely caused by a dominating 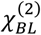 from the lipid bilayer. The field sensitivity is thus the result of an interference term between a SONLO and TONLO interaction of 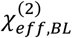. Although the NLO properties of synthetic lipid bilayers have been reported, ^29,30,37^ research that reports and compares *χ*^(2)^ and *χ*^(3)^ for biological membranes is currently lacking due to the enormous complexity of these membranes. Biological membranes differ in lipid composition and possess a full spectrum of integral membrane proteins and membrane related structures which cannot be entirely recapitulated in model systems. Moreover, there is currently no method available to separate *χ*^(2)^ and *χ*^(3)^ of the surface and the asymmetrical lipid bilayer due to the absence of electronic resonances and the limited length scale of the interaction.

Nonresonant homodyne-detected SHG offers a valuable experimental tool for the label-free read-out of charged interfaces.^38^ The heterodyne alternative provides the additional functionality of separating *χ*^(2)^ and *χ*^(3)^ contributions,^23^ however its use is limited to charged interfaces in contact with liquids having low (non-biological) ionic strengths, inducing large Debye lengths and therefore would not be able to extract additional information. Relying mostly on the potential drop inside the membrane, we achieved a detection-efficiency-limited temporal resolution of optical read-out with a sensitivity that is among the highest ever reported, even for the best reported research dyes.^6^ The experimental design is fairly simple as it combines two widely available experimental techniques; a patch clamp rig and a multiphoton microscope. Relying on a widely known, non-excitable cell line (HEK293T), the induced changes in membrane potential can be related to changes in SHG intensity with unprecedented sensitivity (~40% per 100mV) and instantaneous temporal response (at the available temporal resolution) caused by the 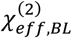 of the lipid membrane. The influence of the SP and interfacial water in cells is more of a static nature, which is moreover corroborated by the increase in SHG intensity upon hyperpolarization, as the total surface electric field of a cell at rest points inwards. Using a voltage clamp configuration on passive cells, the membrane potential can be altered without inducing screening effects, more closely mimicking membrane potential modulation in electrogenic cells and at the same time simplifying the interpretation of the results, offering fundamental insight in the modulation of membrane potential and the application of label-free SHG read-out such as in excitable cells, including neurons and cardiac cells, where TMP is also controlled by a tight control of the ionic imbalance. Previous research which report the voltage sensitivity of SHG imaging using potentiometric dyes, such as the work of Nuryia et al.^4^ report a linear response with a sensitivity of about 10 % per 100 mV. Due to the dominating, resonantly enhanced *χ*^(2)^ of the chromophores, lower excitation powers can be used (1-10 mW). In our experiments, higher powers ranging between 20 and 400 mW are necessary to be able to detect an SHG signal with a sufficient signal-to-noise ratio. It can be reasonably conjectured that, due to the very high SHG and fluorescence signals that are generated by potentiometric dyes, ^6–8^ the presence of any label-free component inherent to the biological membrane (i.e. much lower in intensity but much higher in sensitivity) was never noticed in the past.

We analysed the different components contributing to the SHG response in this complex biological system which is the cell membrane. It seems that the field sensitivity is the result of an interference term between SONLO and TONLO effects within the bilayer. An important conclusion is that the contribution of water molecules at the surface does not generate a dynamic response. Indeed, our experiments are in line with a response which is purely originating from the lipid bilayer. This implies that this label-free tool, monitoring the temporal dynamics of the electric field within the membrane, would offer more detailed information, for example a deeper understanding of how neurons can reach incredible speeds of electric field propagation along their membrane. The temporal resolution of our set-up however, which relies on raster-scanning with a Galvano mirror, is currently limited to a temporal resolution of about 100 ms, 2 orders of magnitude too slow for tracking action potentials in neurons. Temporal limitations on probing purely electronic processes with SHG are imposed by the cell membrane capacitance, collection efficiency of the system, and the time it takes to repetitively scan a region of interest. Nonetheless we foresee that endogenous SHG voltage imaging of electrogenic cells can be achieved by improving the time resolution. This can be accomplished by two main strategies.^39,40^ The most obvious strategy is to increase microscope scanning speed.^41^ The second one is to parallelize the imaging process. Recent advancements include widefield counterpropagating SHG geometries,^42^ lens-free imaging approaches,^43^ harmonic holography,^44^ multi-confocal imaging,^45^ and spatiotemporal widefield illumination.^46^ When using the widefield geometry instead of point scanning, combined with gated detection,^47^ the sensitivity can be increased by 2-3 orders of magnitude, resulting in a temporal resolution of up to 1 ms.^47^

Label-free, NLO interrogation of cellular activity could open up new avenues for understanding cellular function and communication in complex cellular networks, essential for understanding the brain. With the foreseen improvements in temporal resolution, SHG coming from the membrane interface, has the potential to emerge as a powerful non-invasive read-out modality.

## Materials and methods

### HEK293T cell culture

Unless indicated, all compounds were purchased from Life Technologies. Cell lines were maintained in Dulbecco’s Modified Eagle’s Medium without phenol red (Fluobrite) + GlutaMAX (high glucose) supplemented with 10 % Heat Inactivated Fetal Bovine Serum, 1 % non-essential amino acids and 1 % Pen Strep (10,000 units/ml penicillin; 10,000 *μ*g/ml streptomycin stock concentration). Cells were grown in T75 flasks and placed in an incubator (Binder) at 37 °C and 5 % CO_2_. HEK293T cells were grown until they reached 90 % confluency, and split 1/10 two times a week. First, the medium was removed and the cells were gently washed with Dulbecco’s Phosphate Buffered Saline without CaCl2 and MgCl2. Cells were dissociated from the flask by adding 1 ml Versene solution (0.48 mM) for 2 min at 37 °C. The flask was manually agitated and checked under a microscope to make sure all the cells were detached from the surface. New flasks were seeded depending on the starting confluency and 15 ml of prewarmed medium was added. Glass coverslips (25 mm diameter and 0.16 mm thickness) were sterilized by immersion in pure ethanol followed by flaming and placed in six-well plates. A Poly-L-Lysine solution of 0.5 mg/mL was prepared in 150 mM borate buffer (pH 8.5), 2 ml was added to each well and left to incubate for at least 30 min before rinsing the coverslips 3 times with Dulbecco’s Phosphate Buffered Saline for 10 min. 1 ml of seeding solution was added to the coverslips at a final density of 2000 cells/ml (Scepter cell counter, Millipore) and supplemented with 4 ml of medium. They were left to attach for at least 2 hours before the start of the experiments. Prior to electrophysiology and imaging, coverslips were washed with Dulbecco’s Phosphate Buffered Saline and transferred to a custom-made recording chamber. Extracellular fluid was added (see Table 1 for the composition). Osmolarity was adjusted to 320 mOsm by adding glucose (Osmomat-3000-basic, Gonotec, Germany).

**Table 1:**
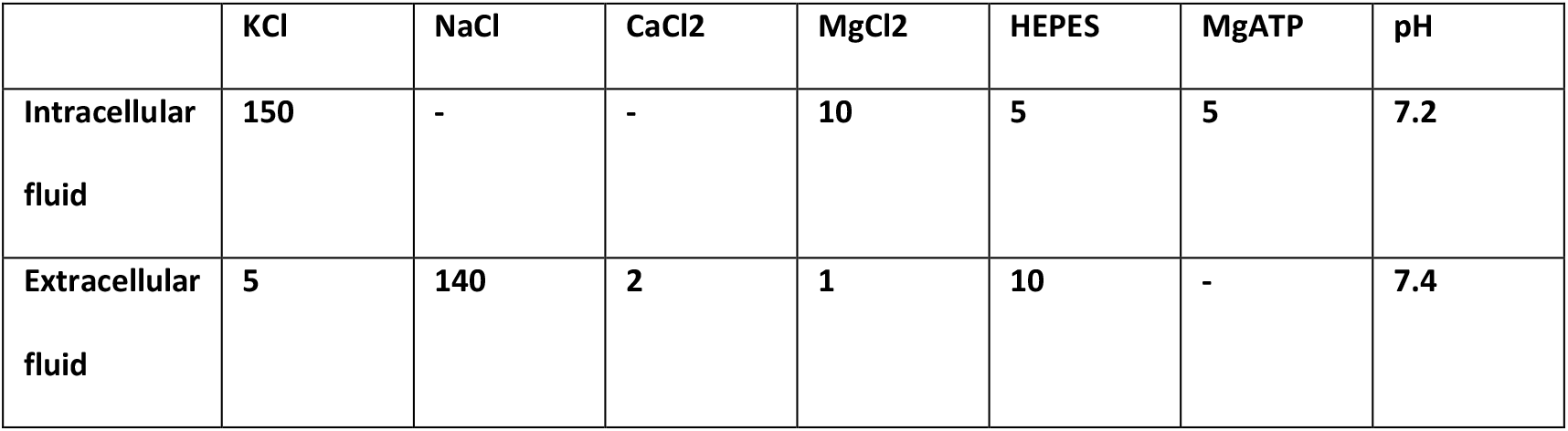
Concentration (in mM) of the different components in intracellular and extracellular medium and their final pH

|  | KCl | NaCl | CaCl<sub>2</sub> | MgCl<sub>2</sub> | HEPES | MgATP | pH |
| --- | --- | --- | --- | --- | --- | --- | --- |
| **Intracellular fluid** | **150** | - | - | **10** | **5** | **5** | **7.2** |
| **Extracellular fluid** | **5** | **140** | **2** | **1** | **10** | - | **7.4** |

### Electro-optical setup

The microscope (BX61WI, Olympus) was coupled to a mode-locked femtosecond laser (Insight DS+, spectra-Physics) producing a vertically polarized beam with an average power output of 1.1 W at 1000 nm and a repetition rate of 80 MHz with pulse widths of 120 fs. The output power was controlled by a combination of a polarizer and achromatic half-wave plate. The laser excitation wavelength for all experiments was set at 900 nm. The microscope allowed to measure both in transmission and reflection with 4 photomultiplier tubes for SHG and TPF using a Galvanometer scanner. Filter cubes consisted of a narrow bandpass filter for the SHG at 450 nm and a 470 nm longpass filter for the TPF. An immersion objective (40X, 0.80 NA, 3.5 mm WD, Nikon) was used. The setup was mounted on an anti-vibration optical table (TMC, USA) and a custom-made Faraday cage surrounding the microscope and setup was made using aluminium and stainless steel. The membrane potential was controlled by clampex software (pClamp, Molecular Devices). The setup consisted of an amplifier (Axopatch 200B-2), an analog to digital converter (Digidata 1550 low-noise data acquisition system) and the micromanipulator (PS-7000C patchStar Micromanipulator) all purchased from Molecular Devices (UK). As a recording electrode, a chlorinated silver wire (0.25 mm in thickness) was used (1-HLA-005, Molecular Devices, California, USA). The electrode was regularly abraded with sandpaper and immersed in commercial bleach solution for about 30 min in order to keep it chlorinated. The reference electrode was a grounded Ag/AgCl pellet electrode (1-HLA-003, Molecular devices, California, USA). Borosilicate glass pipettes with filaments (BF150-86-7.5) were pulled on a P-1000 Flaming/Brown Micropipette Puller (both from Sutter Instruments, California, USA) with an optimized protocol for obtaining 2-5 *MΩ* resistance. Patch pipettes were filled with intracellular fluid using a non-metallic syringe needle (MicroFil, World Precision Instruments, Florida, USA). Intra- and extracellular fluid compositions can be found in Table 1 and the osmolarity was adjusted to 300 mOsm (Osmomat-3000-basic, Gonotec, Germany) by adding glucose. Extracellular fluid was stored at 4 °C and intracellular fluid at −20 °C (aliquoted in Eppendorf tubes). All chemicals were purchased from Sigma-Aldrich and exhibited a purity degree above 95 %.

## Acknowledgments

K.C. and C.B. acknowledge the support of the KUL Research Grant IDO/12/007. C.B. acknowledges the support of the FWO Research Grant G0947.17N and the KUL Research Grant OT/14/084. K.C. acknowledges the support of the FWO Research Grant G0A1817N. T.V. acknowledges financial support from the Hercules Foundation (AKUL/11/15), the KU Leuven (ID-N/19/014), and the Fund for Scientific Research-Flanders (Research Project G099319N). O.D. acknowledges the KU Leuven Research Council for financial support (C14/16/063 OPTIPROBE). S.V.C. acknowledges financial support from the Fund for Scientific Research-Flanders (FWO-V/G099319N) and the KU Leuven (starting Grant/ID-N/19/014).

